# Investigation of the nicotinic receptor EAT-2 as a novel target to mitigate plant parasitic nematode infections

**DOI:** 10.64898/2026.03.20.711559

**Authors:** Henry Atemnkeng Nvenankeng, Elizabeth Hatch, Jefferey R. Thompson, Jim Goodchild, Philippa Harlow, Lindy Holden-Dye, Vincent O’Connor

## Abstract

Plant parasitic nematodes (PPNs) are microscopic soil dwelling pests that infect crops, using a lance-like organ, the stylet, to hatch, invade plant roots, and establish feeding sites. Stylet function is underpinned by pharyngeal muscle contraction and relaxation cycles, making it an attractive route to disrupt the PPN lifecycle. However, knowledge of pharyngeal regulation in PPNs is relatively limited. In the free-living nematode *Caenorhabditis elegans*, the nicotinic receptor EAT-2 stimulates pharyngeal contraction to facilitate feeding. Here we hypothesize that EAT-2 orthologues may regulate a similar function in PPNs. A phylogenetic analysis reveals that EAT-2 and its orthologues in other nematode species cluster as a distinct group suggesting that EAT-2 is exclusive of other animal species. We identified *eat-2* in the genome of the potato cyst nematode *Globodera rostochiensis* and used *in situ* hybridization to establish an anterior expression pattern consistent with a pharyngeal function. *In vitro* pharmacological assays directly compared the response of *C. elegans* pharynx and *G. rostochiensis* stylet to cholinergic compounds. Both pharyngeal and stylet activity were stimulated by acetylcholine and nicotine, and these responses were blocked by the nicotinic receptor antagonists, mecamylamine and tubocurarine. These data are consistent with a conserved cholinergic pathway mediated by EAT-2 regulating pharyngeal muscle function. It highlights EAT-2 as a potential determinant of stylet thrusting and a promising pharmacological target to selectively mitigate PPN infections.

---

Plant parasitic nematodes (PPNs) are global economic pests that account for significant annual losses in agriculture (Lilley et al., 2024). Current control strategies rely on farming practices like crop rotation, fallowing, use of resistant crop varieties and chemical strategies that disrupt the parasitic life cycle (Li et al., 2015; Mesa-Valle et al., 2020; Pires et al., 2022). Unfortunately, several available nematicides are either restricted or banned because of their broad actions and negative environmental impacts (Ngala et al., 2021). Thus, finding novel pesticide targets and new products that meet required regulatory standards is of high importance (EPA, 2023). Generally, the nervous system of pests is an attractive target for pesticides because they are comprised of several targets that are rapidly and readily druggable (Hirata, 2016). Pest control has been well served by targeting ion channels whose disruption impairs signal transduction resulting in muscle paralysis, spasms and death of invertebrate pests (Crisford et al., 2015; Hirata, 2016; Raisch and Raunser, 2023). Although not widely used, integrating the biological understanding of mechanisms unique to behaviors that support parasitism might refine the focus on pesticide selectivity and reduce the limiting negative environmental impacts.

Investigating the molecular determinants of core behaviors in PPNs is challenging given the lack of genetic tools. In this regard, studies in the model nematode *Caenorhabditis elegans* can be informative (Holden-Dye and Walker, 2014; Coke et al., 2024). The nervous system receives several sensory modalities that co-ordinate and generate distinct phenotypic and behavioral outputs which regulate essential processes. This means that behaviors like feeding, locomotion and reproduction can be templated in *C. elegans* and translated to other nematodes (Alcedo et al., 2013; Chew et al., 2013; Gjorgjieva et al., 2014; Thapliyal and Babu, 2018). The pharynx of *C. elegans* acts as a neuromuscular pump that regulates ingestion of bacteria. In *C. elegans* 20 radial muscle cells organized into 8 pharyngeal muscles regulate feeding in a two-part process involving pharyngeal pumping and peristalsis. The rate and pattern of pharyngeal pumping is controlled by the pharyngeal nervous system (McKay et al., 2004; Avery and You, 2012). The presence of food triggers serotonergic neurons to release 5-HT resulting in elevated pharyngeal pumping rates (Avery and Horvitz, 1990; Song et al., 2013). The biogenic amine 5-HT acts on cholinergic neurons MC and M4 to induce essential muscle contractions that are necessary for feeding (Niacaris and Avery, 2003; Ishita et al., 2020). A core determinant for this response is EAT-2, the receptor that utilizes ACh released from the MC neuron to initiate pharyngeal muscle contraction that opens the pharyngeal lumen to ingest bacteria (McKay et al., 2004).

Interestingly, EAT-2 was initially classified as a non-alpha nicotinic receptor subunit (McKay et al., 2004) because it is missing the signature vicinal cysteines amino acid residues known to be a major molecular determinant of agonist acetylcholine binding (Kao and Karlin, 1986; Blum et al., 2011). Surprisingly, despite lacking this motif, both the *C. elegans* EAT-2 and its orthologue from the animal parasitic nematode *Ascaris suum* have been shown to function as homo-oligomeric receptors made up of the co-assembly of five identical EAT-2 receptor subunits (Choudhary et al., 2020). Moreover, functional expression of EAT-2 requires co-expression with EAT-18, a single transmembrane domain protein that functions as an auxiliary subunit. Intriguingly, and of relevance with respect to potential as a target for selective nematicides, EAT-18 has no known homology to previously described proteins (McKay et al., 2004). EAT-18’s unique and essential contribution is proposed to be via an association with the EAT-2 pentamer at the muscle plasma membrane (Choudhary et al., 2020).

In PPNs pharyngeal function controls critical behaviors using a specialized pharyngeal structure called the stylet, a lance-like hollow structure important for their lifestyle. The neurobiology underpinning this pharyngeal function is poorly understood (Grundler and Böckenhoff, 1997). In one modality pharyngeal muscles co-ordinate the protraction and retraction of the stylet by J2s that it used to pierce the encasing eggshell thus driving hatching (Perry and Clarke, 2000; Mkandawire et al., 2022). This biology is later used by the hatched J2s to pierce plant roots and facilitate their invasion of host plants (Bernard et al., 2017; Pulavarty et al., 2021). Additionally, a distinct pharyngeal function involving stylet thrusting and median bulb pumping allows protruded stylets to ingest nutrient or release effectors and cell-wall degrading enzymes into feeding sites of invaded host plants (Koga, 2014).

Despite the difference in the feeding style and habits of *C. elegans* and *G. rostochiensis*, the organization of their pharyngeal muscles includes the corpus, isthmus and terminal bulb. Furthermore, the exposure of the whole organism to 5-HT drives pharyngeal pumping responses from averages of about 60 pumps/minute to 250 pumps/minute in *C. elegans* and stylet thrusting behaviors from about zero to 80 thrusts/minute in *Globodera* (Masler, 2007; Song et al., 2013). A putative model based on the limited comparisons of pharyngeal structure and pharmacological evidence suggests that serotonergic signaling regulates pharyngeal function upstream of EAT-2 (Kearn et al., 2017; Ishita et al., 2020). Based on this information, we have investigated the molecular details of EAT-2 in PPNs, its cellular expression and developed pharmacological assays to resolve cholinergic function. The data supports the notion that cholinergic modulation of pharyngeal function via EAT-2 underpins stylet function. This provokes the idea that EAT-2’s function and its distinct molecular features make it a suitable target for new nematicide development.

## MATERIALS AND METHODS

### C. elegans maintenance

*C. elegans,* N2 (Bristol strain) were obtained from the *Caenorhabditis* Genetics Centre (CGC), grown and maintained under standard conditions (Brenner, 1974) and used as wild type (WT) worms.

### G. rostochiensis maintenance

Infective juveniles (J2s) of *G. rostochiensis* were hatched by incubating nematode cysts in potato root diffusate (PRD) over 7 days at room temperature (Gaihre et al., 2019). Freshly hatched J2s (1 day old) were washed in M9 buffer containing 0.01% w/v Bovine Serum Albumin (BSA) prior to experimentation. Batch hatchings were performed for each experiment to obtain freshly hatched juveniles. Unused worms were collected in batches and stored at -20 °C before being used for RNA extraction. Nematode cysts used were from the James Hutton Research Institute, Scotland and provided by Vivian C. Blok. The PRD used for hatching was provided by Catherine Lilley, University of Leeds, UK.

### Drugs and chemicals

Serotonin creatinine sulphate monohydrate, Nicotine hydrogen tartrate salt, Mecamylamine hydrochloride and Tubocurarine chloride were purchased from Sigma Aldrich, UK. To make stock solutions that were used in experiments the drugs were dissolved in sterile ddH2O and stored at -20 °C. These were used no longer than 1 month after preparation.

### Characterizing C. elegans and G. rostochiensis response to 5-HT, ACh and nicotine

#### C. elegans behavioral assay

These assays were performed on solid NGM plates by supplementing molten NGM with 5-HT, nicotine or ACh to achieve the indicated final concentration. A *C. elegans* pharyngeal pump dose-response to 5-HT was established by testing concentrations up to 5mM. ACh and nicotine were tested at 10 mM. 3 ml NGM containing vehicle or the indicated concentration of drug was transferred into clear Corning Coster 6-well plates. After setting the NGM plates were stored at 4 °C and used the following morning. Before experimenting, these plates were equilibrated to room temperature for 30 minutes. Well-fed young adult *C. elegans* (L4+1s) were washed in M9 buffer to remove bacteria and then picked onto test plates. Pharyngeal pump rates were measured by making visual observations using a Nikon SMZ800 binocular microscope. A pump was defined as a forward and backward movement of the grinder and was recorded for a minute per worm (Raizen et al., 1995). All *C. elegans* behavioral assays were performed on NGM supplemented plate unless specified otherwise.

### G. rostochiensis behavioral assay

Stylet thrusting dose-response to 5-HT was established by supplementing 20 mM HEPES solution, pH 7.4 with indicated concentrations of 5-HT. Ten freshly hatched J2s washed 3 times with ddH2O were pipetted in 2 µl and transferred into clear Corning Coster 96-well plates containing 200 μl of 20 mM HEPES (control) or with an indicated 5-HT concentration. The number of stylet thrusts were measured for 30 seconds per worm, 30 minutes after incubating G*. rostochiensis* in 5-HT and represented as number of thrusts per minute. A forward and backward movement (extension and retraction) of the stylet was counted as one stylet thrust (Kearn et al., 2017). All *G. rostochiensis* behavioral assays were performed in liquid assays unless specified otherwise.

### Acute responses to selected drugs

We investigated the acute effects of 10 mM nicotine, 200 μM mecamylamine and 300 μM tubocurarine on 5-HT induced pharyngeal pumping and stylet thrusting. This was performed by adding respective drug or requisite vehicle after establishing a steady-state stylet thrusting response for 30 mins in 1mM 5-HT. The acute response to these selected drugs on 5-HT induced pharyngeal pumping and stylet thrusting was measured for up to 2 hours

### Chronic responses to selected drugs

The chronic effect of the drugs was investigated by pre-incubating *C. elegans* or *G. rostochiensis* for 24 hours in 10 mM nicotine, 200 μM mecamylamine and 300 μM tubocurarine prior to transferring them onto assays plates.

### Molecular biology

#### Phylogenetic analysis

We used the basic alignment search tool (BLAST) (Altschul et al., 1990) and the query *Ce*.EAT-2 (Uniprot accession number: Q9U298) to search for EAT-2 homologues on Uniprot and WormBase ParaSite. The resulting putative protein sequences were imported into the MEGAx (Molecular Evolutionary Genetics Analysis) tool and aligned via MUSCLE alignment (multiple sequence alignment by log-expectation) (Stecher et al., 2020) with default parameters. The aligned sequences were used to generate a phylogenetic tree using the Maximum Likelihood (ML) statistical method, a Jones-Taylor-Thornton (JTT) substitution model and 500 bootstrap replications to assess node support. The tree interference options were set to default parameters. The phylogenetic tree output was exported in a Newick format and visualized using iTOL (interactive tree of life) (Letunic and Bork, 2024). The following protein sequences were used in the analysis to build the phylogenetic tree:

Q9U298 (*C. elegans*); A0A261C1U6 (*C. latens*); A0A9P1MVH6 (*C. angaria*); A0A158PA63 (*A. cantonensis*); A0A0N4XX74 (*N. brasiliensis*) A0AA36C8X6 (*M. spiculigera*); A0AAF3EAP6 (*M. belari*); A0AA39GWV7 (*S. hermaphroditum*); A0A7E4V0M4 (*P. redivivus*); A0A2A6C485 (*P. pacificus*); A0A0K0DZ20 (*S. stercoralis*); A0A811LPF4 (*B. okinawaensis*); A0A1I7S0H9 (*B. xylophilus*); A0A914M6R1 (*M. incognita*); A0A6V7VQB4 (*M. enterolobii*); A0A914I8V5 (*G. rostochiensis*); A0A8T0A120 (*M. graminicola*); A0A914XFN7 (*P. sambesii*); A0A0N4UD55 (*D. medinensis*); A0A0B2W3Z2 (*T. canis*); A0A4E9FW31 (*B. malayi*); W2TG30 (*N. americanus*); A0A4U8V1R9 (*S. carpocapsae*); A0A2G9V6F1 (*T. circumcincta*); A0AA36MEL8 (*C. nassatus*); A0A0N4VD61 (*E. vermicularis*); A0A1I7V6L7 (*L. loa*); A0A0V0XKJ1 (*T. pseudospinralis*); A0A9C6WDJ3 (*D. albomicans*); A0A8B8HQB2 (*V. tameamea*); M9PFD8 (*D. melanogaster*); A5H031 (*M. domestica*); A0A9P0BVQ3 (*C. includens*); A0A8R2AB14 (*A. pisum*); A0A8B7PB12 (*H. azteca*); A0A8I6SIA1 (*C. lectularius*); AgR003_g027 (*A. suum*); A0A915M3T9 (*M. javanica*); tig00002135.g43658.t1 (*M. arenaria*); A0A1I8BSG2 (*M. hapla*); Hsc_gene_20958.t1 (*H. schachtii*), A0A183C532 (*G. pallida*); KAH7704458.1 (*A. avenae*); A0A158PA63 (*H. sapiens*).

### RNA extraction

Total RNA was extracted from one 1000 *C. elegans* young adults and 2000 infective *G. rostochiensis* juveniles respectively. Nematodes were washed with diethyl pyrocarbonate (DEPC) treated M9 buffer, centrifuged at 500 g and the supernatant discarded and leaving the worms in minimal volumes of DEPC in the 2 ml eppendorf tubes. 1 ml of TRIzol^TM^ was added to each sample and homogenized for a minute using the hand-held VWR® VDI 12 homogenizer (VWR Collection). Homogenization was monitored via visual inspection and was continued until all intact organisms were broken open. Total RNA was isolated using the sequential TRIzol^TM^ (Invitrogen) Qiagen RNA clean up protocol (Metzger, 2024). Total RNA extracted was stored at -20 °C or used to synthesize cDNA using the SuperScript® III First-Strand Synthesis System protocol (Invitrogen).

### PCR amplification

The sequences that encode the open reading frame (ORF) for *Ce*.EAT-2 (Uniprot accession number: Q9U298) was used as input to perform a protein BLAST search on Uniprot and WormBase ParaSite. We identified orthologues in sequenced parasitic nematodes and the predicted ORF for *Gr.*EAT-2 (accession number: A0A914I8V5). Primers designed and synthesized with Integrated DNA Technologies were used to amplify the predicted coding sequence of both organisms. The following primers were used to amplify EAT-2 transcripts for *C. elegans* and *G. rostochiens;*

*FwCe.eat-2* 5’-CGCAC***<u>gaattc</u>***ATGACCTTGAAAATCGCATT-3’

*RvCe.eat-2* 5’-CAGGCTAACAACTATAACTTATT***<u>cccggg</u>***CG- 3’

*FwGr.eat-2* 5’-GC***<u>ggtacc</u>***ATGTTTTTGCGA-3’

*Rv.Gr.eat-2* 5’-GGTGGAGAGATT***<u>cccggg</u>***GA-3’

Restriction sites (underlined sequences) were added at primer ends to flank the amplified sequence. The authenticity of the amplified sequence was confirmed by sequencing the forward and reverse strands of the amplified gene (Eurofins Genomics) (see supplementary 1).

### Hybridization Chain Reaction RNA-Fluorescence in-situ hybridization

The expression patterns for the target mRNA *Gr.*EAT-2 (Uniprot accession number: A0A914I8V5), *Gr.*MYO-3 (Uniprot accession number: A0A914HJT9) and *Gr*.UNC-17 (Uniprot accession number: A0A914GTZ9) were investigated using a modified HCR^TM^ RNA- FISH (v3.0) protocol for whole-mount nematode larvae (Choi et al., 2016).

### Probe sets, amplifiers and buffers

Probes against target transcripts, probe hybridization buffers, probe wash and amplifier buffers were purchased from Molecular Instruments. The number of probe binding sites for *Gr.eat-2* was set to 40 to mitigate likely low and restricted expression levels based on exemplar data in *C. elegans* (McKay et al., 2004). Probe binding sites for *Gr.unc-17* and *Gr.myo-3* were standard 20. Amplifiers and hairpins used were: B1H1, B1H2 for EAT-2 and B2H, B2H2 for UNC-17 and MYO-3. Other reagents used included: M9 buffer (22 mM KH2PO4, 42 mM Na2HPO4, 20.5 mM NaCl, 1 mM MgSO4), PBST (1X PBS, 0.1% Tween), 4% paraformaldehyde (PFA), glycine solution (2 mg/mL glycine, PBST), proteinase K solution (100μg/mL).

### Preparation of fixed whole-mount nematode larvae

Approximately 1000 *G. rostochiensis* J2s were washed three times with 1 mL of M9. During each wash the tube were centrifuged at 500 g for 2 min to bring larvae to the bottom and the supernatant carefully removed. Washed J2s were aliquoted in minimal volumes of M9 buffer and incubated in 1 mL of 4% paraformaldehyde (PFA) before immediately freezing at -80 ^◦^C overnight. The following day the fixed J2s were thawed at room temperature for 45 minutes washed twice in 1 mL of PBST and treated with 1 mL proteinase K (100 µg/mL) (Sigma Merck) for 10 min at 37 ^◦^C. The proteinase K treated J2s were washed with PBST and incubated in 1 mL of glycine solution (2 mg/mL) for 15 min on ice followed by 2 washes with PBST (HCR^TM^ RNA-FISH (v3.0) protocol).

### Probe hybridization, amplification and detection

Fixed and permeabilized J2s were incubated in 1 mL of 50% PBST / 50% probe hybridization buffer for 5 min at room temperature and centrifuged at 500 g for 2 min before removing the supernatant. These processed J2s were then pre-hybridized in 300 µL of HCR™ probe hybridization buffer at 37 ^◦^C for 1 h. The indicated probe solutions were prepared just before use by adding 2 µL of the 1 µM stock to 200 µL of probe hybridization buffer at 37 ^◦^C (See the HCR^TM^ RNA-FISH (v3.0) protocol for more detail) (Choi et al., 2016). The pre-hybridized J2s were added to the freshly made probe solution and incubated overnight (>12 h) at 37 ^◦^C. After the overnight incubation the probe solution was removed by washing the J2s 4 times in 1 mL of probe wash buffer at 37 ^◦^C for 15 minutes. These washes were followed by two further 5 mins washes at room temperature in 1 mL of 5× SSCT. To facilitate pelleting of worms between washes prepared worms were centrifuged at 800 g. Hybridized J2s were incubated in 300 µL of amplification buffer for 30 min before adding 200 µL of the hairpin solution and incubating overnight (>12 h) in the dark, at room temperature. Following this incubation the worms were washed 5 times with 1 mL 5× SSCT at room temperature. These samples were treated with 0.01% DAPI (v/v) and stored at 4 ^◦^C protected from light before imaging.

### Mounting and imaging

The probe treated specimens were pipetted in 20 µL of SSCT (about 30-50 nematodes), placed on a glass slide, secured with a cover slip and sealed with nail varnish. Slides were mounted on the Olympus/Yokogawa Spinning Disk confocal microscope and imaged using the Alexa fluor 488 for GFP and 594 for mcherry. Images were prepared using image J, with minor adjustments on the brightness and contrast of the DIC images (Schindelin et al., 2012).

### Statistical analysis

Data analysis was performed with GraphPad Prism 10.4 and presented as the mean ± standard error of the mean (SEM) for a number of observations (n). Statistical significance was tested either by unpaired student’s t-test, one-way or two-way ANOVA followed by Bonferroni multiple comparison where appropriate. Significance levels were set at p<0.05. Every experiment was repeated in 3 or more independent occasions, unless stated otherwise. EC_50_ values with 95% confidence intervals were determined by plotting log (agonist) vs. normalized response-variable slope and fitted to the equation; Y=100/(1+10^((LogEC_50_-X)*HillSlope))

## RESULTS

### Phylogenetic evidence reveals EAT-2 unique expression in the Nematoda

Amino acid sequence for *C. elegans* EAT-2 was used as a query to run a protein blast on Uniprot and WormBase ParaSite databases. The phylogenetic analysis revealed *Ce.*EAT-2 and orthologues in other nematode species cluster as a distinct group suggesting that EAT-2 orthologues are exclusive of other animal species (**Fig. 1A**). The EAT-2 orthologues predict the classical nicotinic acetylcholine receptor transmembrane topology encompassing a large extracellular N-terminal domain and four intervening transmembrane helices leading to a short extracellular C-terminus. In the context of a nAChR the acetylcholine binding site lies at the interface between the extended N-terminal extracellular domains of neighboring subunits. In homomeric receptors like alpha-7 and EAT-2 this is at the principal (front) and complementary (back) end of each subunit. The six-loop fold that comes together to contact the activating ACh has a major and minor contact made up of discontinuous loops ABC and DEF respectively. Comparing amino acid residues that make up the principal and complementary phases of the ligand binding domain for *Ce.*EAT-2 and its closely related human orthologue the nicotinic receptor alpha-7, revealed a 50% conservation of amino acid residues in these loops (**Fig. 1B**). A multiple sequence alignment for *Ce*.EAT-2 and some selected nematode species with alpha- 7 revealed the absence of vicinal cysteine residues in the loop-C region of these receptor subunits (**Fig. 1C**). Interestingly these pivotal vicinal cysteines were originally identified as an absolute requirement for nicotinic receptor alpha subunits acetylcholine binding, but emerging data recognizes that activation can occur in the absence of this motif (Blount and Merlie, 1990; Blum et al., 2011; Choudhary et al., 2020).

**Fig. 1.**
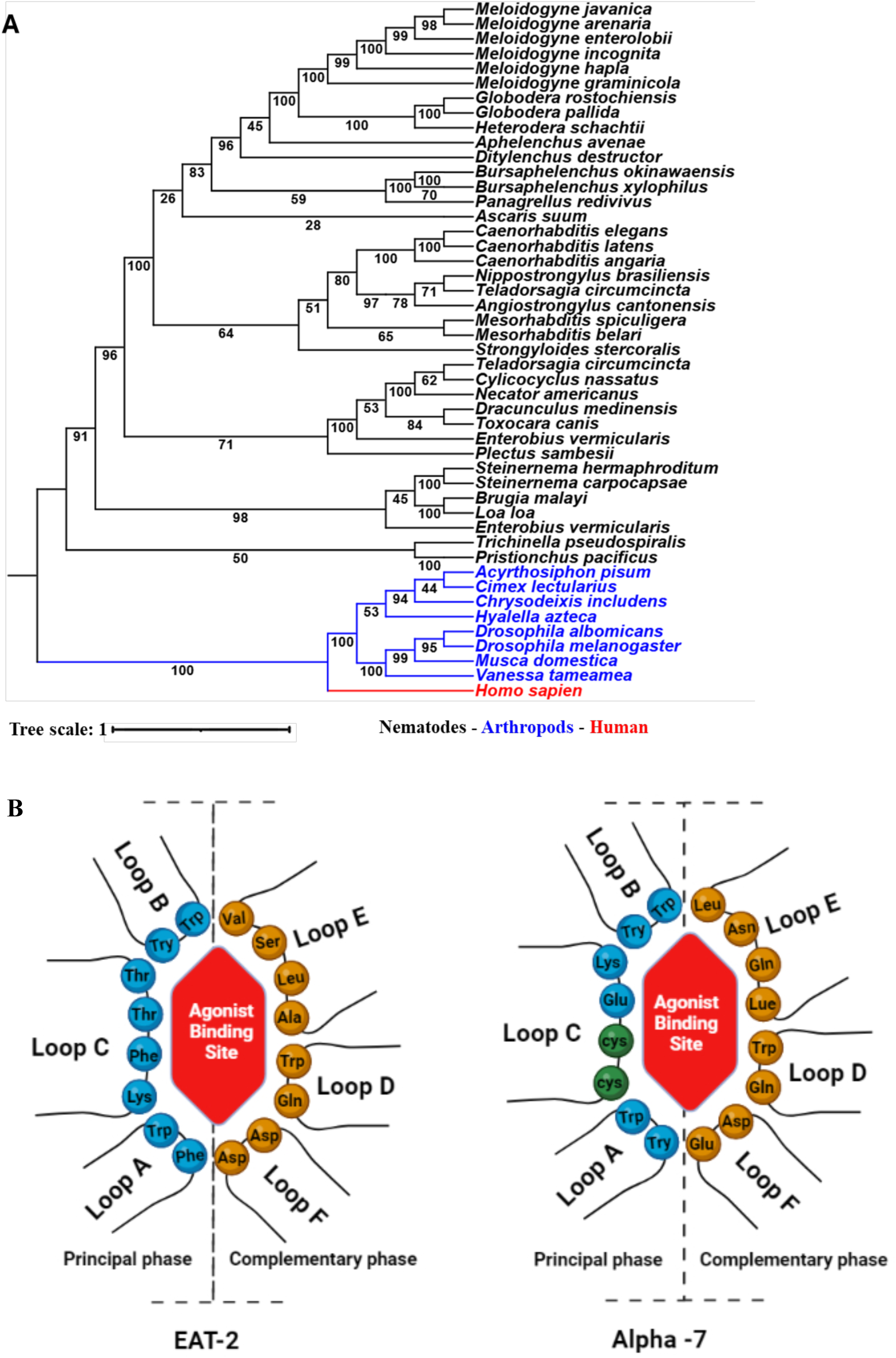
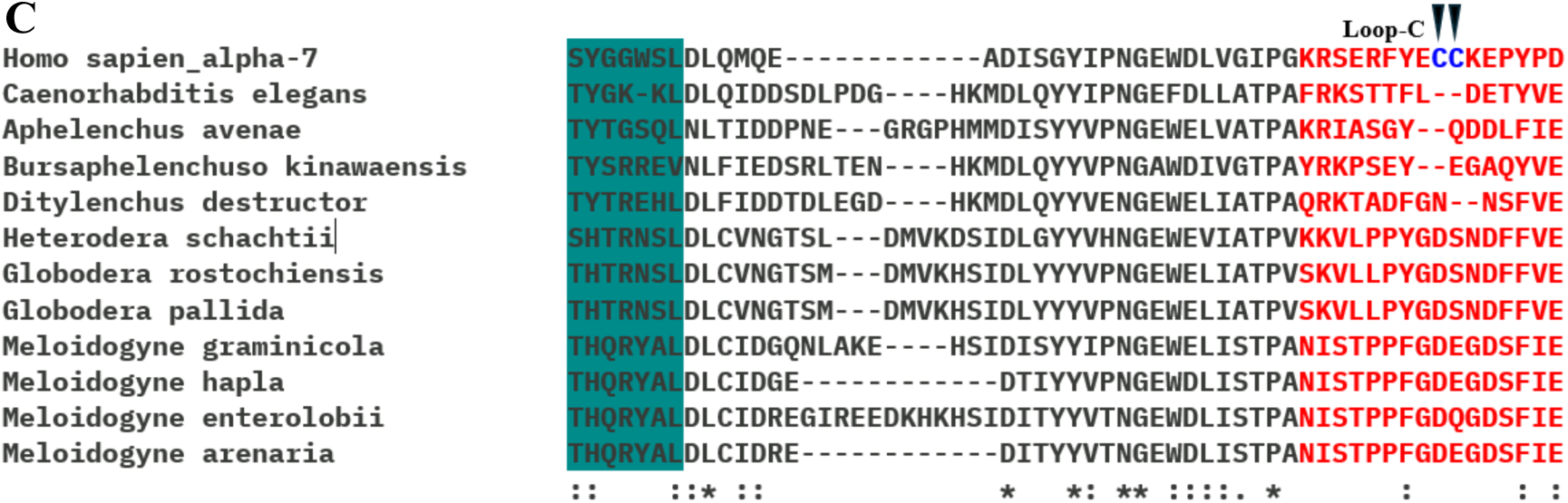
Maximum likelihood phylogeny of EAT-2. **A.** The tree was rooted with α-7, the closest human orthologue of *C. elegans* EAT-2. Phylogenetic analysis groups EAT-2 of nematodes (green) in a common branch separate from orthologues of arthropod and humans. **B.** A schematic representation showing a 50% conservation of amino acid residues that make the agonist binding site (ABS) of EAT-2 and α-7. Loops A, B and C make up the principal phase of the binding domain and loops D, E and F, the complementary phase. **C.** A multiple sequence alignment for *C. elegans* EAT-2 and orthologues in some economically important plant parasitic nematodes shows the absence of vicinal cysteines in the loop-c when compared to the human α-7.

### EAT-2 expression pattern in G. rostochiensis

#### Spatial expression of Gr.EAT-2 HCR RNA-FISH

To resolve the organization of cholinergic transmission and the molecular features of *Gr*.*eat*-2 we first authenticated the predicted sequence. This involved extracting total RNA from *G. rostochiensis,* reverse transcribing it to cDNA and then using specific primers to amplify the predicted ORF for *Gr.eat-2.* The amplification product was authenticated through sequencing and fully aligned with the predicted sequence. The amplified sequence was used to design probes for HCR RNA-FISH, to investigate the spatial expression of *Gr*.*eat-2* transcripts using whole-mount in-situ hybridization. We extended this to the wider cholinergic system using the predicted orthologues for *Gr.unc-17* which encodes the vesicular ACh transporter in *C. elegans* (Alfonso et al., 1993) and is found in all cholinergic releasing neurons and *Gr.myo-3* which encodes the myosin heavy chain protein highly expressed in longitudinal muscles of the body wall muscle (Meissner et al., 2009). Previously developed methods for getting small molecules like DAPI into worms suggested the need to physically break open the cuticle (Lilley et al., 2018). However, we used the HCR^TM^ RNA-FISH (v3.0) protocol for whole-mount nematode larvae (Choi et al., 2016) and achieved a 30% success rate (n=100). With probes for *Gr*.*eat-2* and *Gr*.*unc-17* in a co-localization hybridization reaction we identified a discrete expression pattern for *Gr*.EAT-2 in pharyngeal muscles, specifically in the median bulb and further discrete but not always present staining in what looked to be the pharyngeal glands (**Fig. 2A**). *Gr*.UNC-17 was identified as punctate staining in the anterior region and along the ventral side of the worm, highlighting the possible locations of cholinergic motor neurons (**Fig. 2A**). The specific signals in the pharynx are suggested to highlight the neuronal cell bodies of the cholinergic neurons that innervate pharyngeal muscles. The specificity of *Gr*.EAT-2’s pharyngeal expression pattern was reinforced in a co-localization hybridization reaction with *Gr*.MYO-3 (**Fig. 2B**). Worms fixed and prepared in the absence of target probes showed no signal for our target genes. The sclerotized head and stylet were auto fluorescent and very bright for both control and probe treated specimen (**Fig. 2C**).

**Fig. 2.**
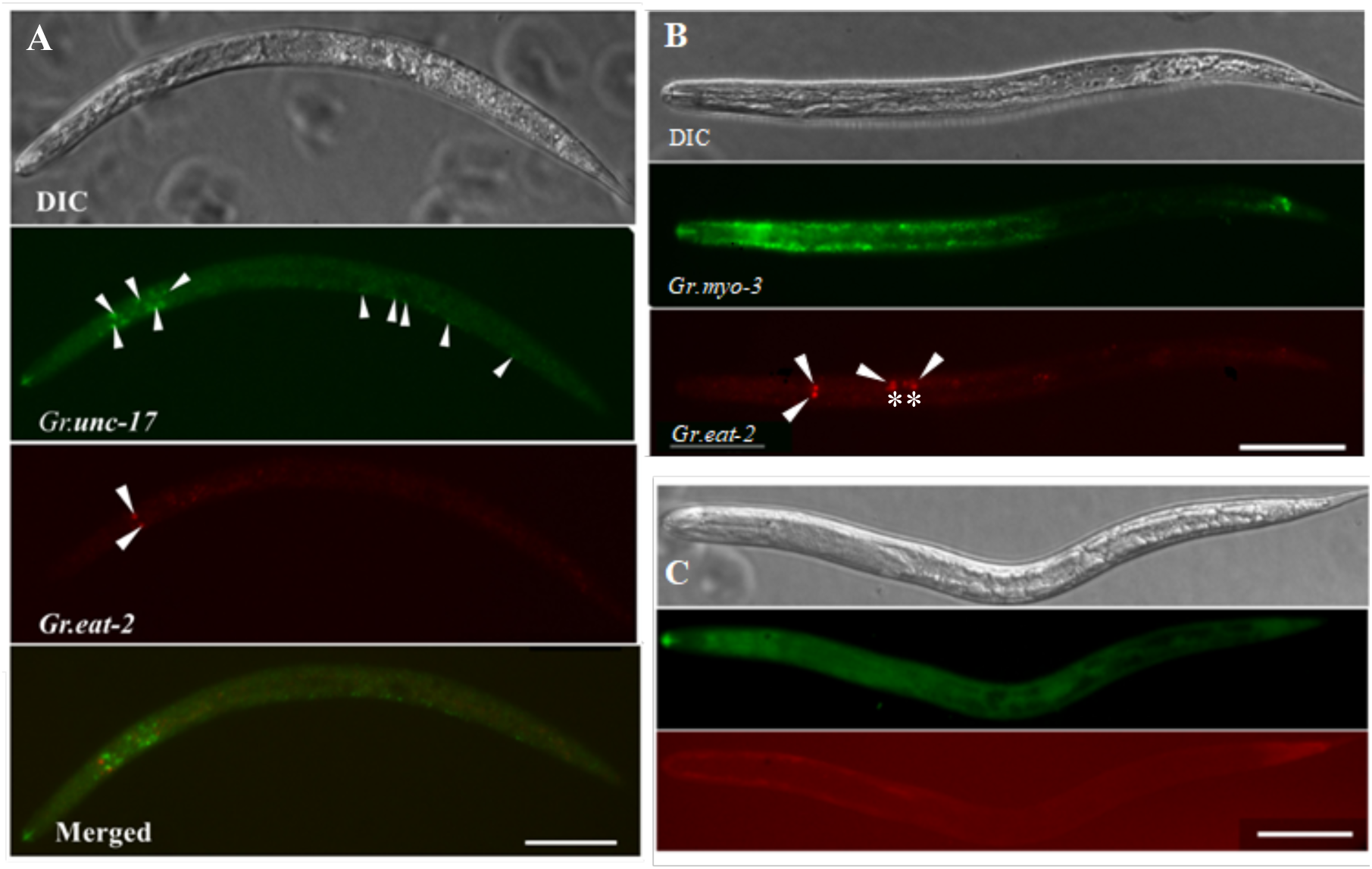
Spatial expression of EAT-2 in *G. rostochiensis* visualized with FISH. **A.** Whole-mount nematode visualizing the expression of *Gr*.EAT-2, seen as red puncta and *Gr*-UNC-17 seen as green puncta in the anterior region, consistent with pharyngeal expression. **B.** *Gr.*MYO- 3 expressed in longitudinal muscles of the body wall muscle (green), **C.** Worms treated without target probe show no cellular expression but a non-specific autofluorescence in the head region and the stylet. Arrow heads point to the localization of indicated hybridization probe; the asterisk represents an additional puncta pattern not always observed in worms that show the arrow heads. The images are representative images from 2k prepared J2s in two independent histology runs. Scale bar, 50 μm.

### A comparative pharmacological analysis of C. elegans pharyngeal pumping and G. rostochiensis stylet thrusting

The functional control of the nematode pharynx is regulated by modulatory neurons that can be probed by the external exposure to 5-HT (Hobson et al., 2006; Horvitz et al., 1982; Perry et al., 2004). We compared the dose-dependence of 5-HT on *C. elegans* pharyngeal pumping and *G. rostochiensis* stylet thrusting to benchmark pharmacological comparisons between these nematode species. Increasing concentrations of 5-HT induced increased pharyngeal activity for both nematode species. In *C. elegans* 5-HT had an EC^50^ of 237 μM (95% confidence limit 168.5 - 315 μM) and 5 mM induced the highest pharyngeal pumping rate. Similarly, in *G. rostochiensis* increasing 5-HT concentrations induced increasing stylet thrusting responses with an EC_50_ of 409 μM (95% confidence limit 365.0 to 457.5 μM) and a maximal stylet thrusting rate at 1mM.

In solutions like M9 or 20 mM HEPES buffer, *C. elegans* moves by thrashing in a coordinated manner (Buckingham and Sattelle, 2009). In contrast, movement in *G. rostochiensis* is relatively uncoordinated. Interestingly, when incubated in 1 mM 5-HT beyond 30 minutes they became immobilized. This was similarly observed by incubating *C. elegans* in 30 mM 5-HT (Ranganathan et al., 2000). *C. elegans* assumed a steady position with elevated pharyngeal pump rates while *G. rostochiensis* adopted a kink position that was associated with the induced stylet thrusting.

We also investigated known agonists of cholinergic systems, ACh and nicotine and observed that these ligands activated pharyngeal pumping and stylet thrusting behaviors. The responses were proportionately higher in *C. elegans* compared to *G. rostochiensis* (**Fig. 3**). The exposure of *C. elegans* to 5-HT induced pharyngeal pumping in similar rates like ACh and nicotine (**Fig. 3A**). With *G. rostochiensis*, the stylet thrusting response to ACh and nicotine was weaker compared to 5-HT induced responses (**Fig. 3B**). In general, the modulation of pharyngeal function by nicotinic receptor agonists supports an underlying contribution of cholinergic transmission within the pharyngeal system of nematodes.

**Fig. 3.**
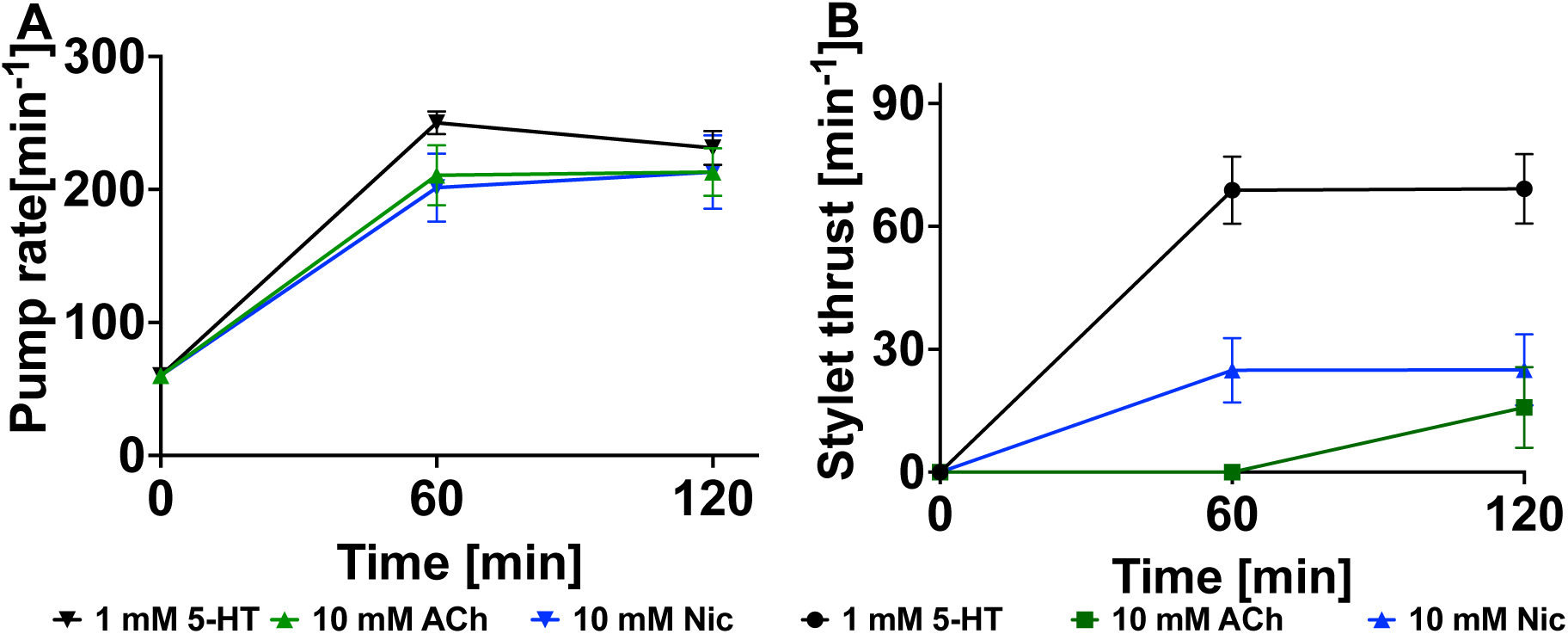
EAT-2 agonists induce pharyngeal behaviors in *C. elegans* and *G. rostochiensis*. **A.** 5-HT, ACh and nicotine activate pharyngeal pump responses in *C. elegans* (n>8) and **B.** stylet thrusting in *G. rostochiensis* (n=10). Data represented as mean ± SEM

### EAT-2 modulators inhibit 5-HT induced pharyngeal function in C. elegans and G. rostochiensis

There is a limited pharmacology of the EAT-2 receptor. However, micromolar concentrations of the agonist nicotine and the antagonists mecamylamine and tubocurarine are reported to be highly potent against the recombinantly expressed EAT-2 receptor (Choudhary et al., 2020). Having established the 5-HT dependence of stylet thrusting and pharyngeal pumping, we performed comparative pharmacological assays to inform us on the organization and role of EAT-2 in the pharyngeal system of PPNs. This involved comparing the acute (short-term) and chronic (long-term) exposure effects of these drugs on 5-HT stimulated pharyngeal responses in both nematode species.

### Acute exposure to cholinergic signaling modulators impact nematode pharyngeal function

The short-term exposure of *C. elegans* to nicotine rapidly inhibited the pharyngeal pump response induced by 5-HT. This inhibitory effect was sustained over the 2-hour experimental period (**Fig. *4*A**). This was an interesting effect because by itself, nicotine activated pharyngeal pumping (**Fig. 3A**) but in combination with 5-HT its effect was rather inhibitory. The presence of antagonists, mecamylamine and tubocurarine induced varied responses on 5-HT stimulated worms. Mecamylamine had a sustained inhibitory effect on 5- HT stimulated pharyngeal pumping in *C. elegans*. In contrast tubocurarine had no effect on this pharyngeal response (**Fig. *4*A)**.

**Fig. 4.**
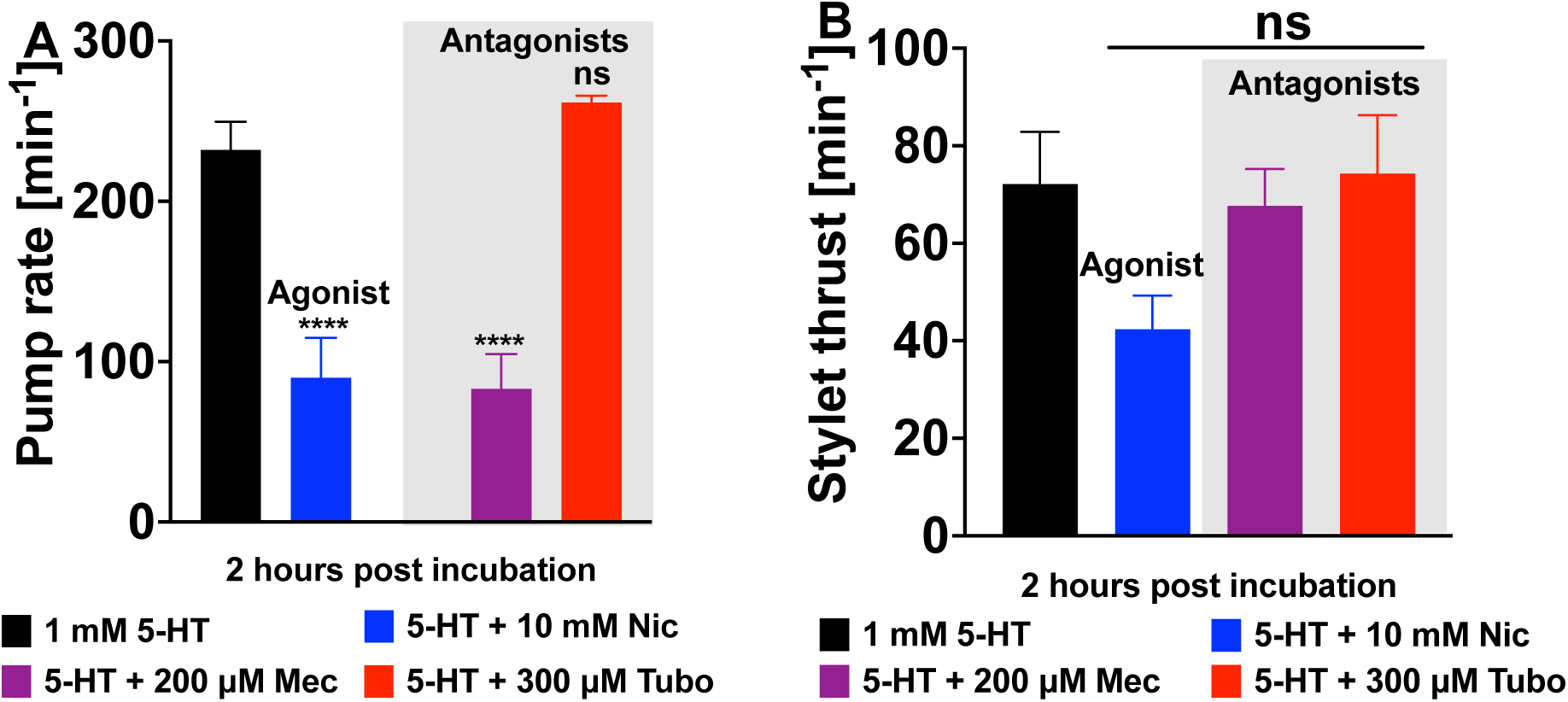
Modulation of pharyngeal function in 5-HT stimulated *C. elegans* and *G. rostochiensis*. **A.** 5-HT induced pharyngeal pump rates in *C. elegans* were inhibited by nicotine and mecamylamine. Tubocurarine had no inhibitory effect, n=10 **B.** In *G. rostochiensis*, these modulators had no significant inhibitory effects on 5-HT induced stylet thrusting, n=13. Data represented as mean ± SEM. The analysis uses the steady state rate of 5-HT as control or 5-HT + ligand and significance tested by two-way ANOVA with Bonferroni’s multiple comparison (^ns^p>0.05, ****p<0.0001).

With *G. rostochiensis,* short-term exposures to nicotine, mecamylamine and tubocurarine failed to significantly inhibit ongoing 5-HT stimulated stylet thrusting. However, although not significant, in the presence of nicotine there was an observable reduction in stylet thrusting (**Fig. *4*B)**

### Chronic exposure to cholinergic signaling modulators impact nematode pharyngeal function

A challenge with nematode drug exposure experiments is the cuticle that restricts the entry and rate of accumulation of drug inside the worm. To overcome this limitation, we prolonged the incubation time of *C. elegans* and *G. rostochiensis* in nicotine, mecamylamine or tubocurarine prior to 5-HT exposure and saw an increase in drug effect.

Pre-incubating *C. elegans* for 24 h in nicotine markedly improved the drugs inhibitory effect on 5-HT stimulated pharyngeal pumping. Similarly, a prolonged incubation in mecamylamine also reduced 5-HT induced pump rates. In contrast prolonged incubations of *C. elegans* in tubocurarine had no effect on the subsequent ability of 5-HT to induce pharyngeal pumping (**Fig. 5A**).

**Fig. 5.**
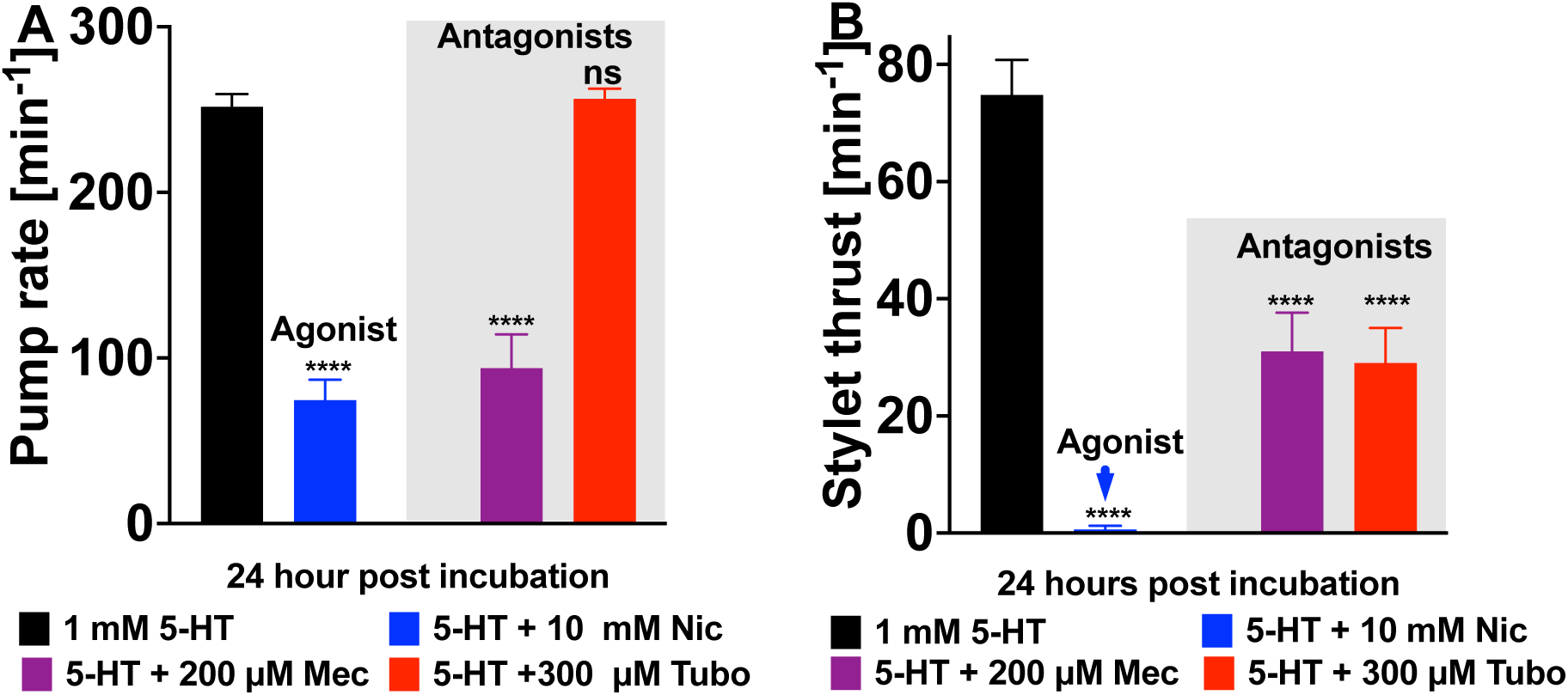
Chronic exposures to cholinergic modulators improves their inhibitory effects on 5-HT induced pharyngeal behaviors in *C. elegans* and *G. rostochiensis*. **A.** Prolonged incubations of *C. elegans* in 10 mM nicotine and 200 μM mecamylamine prior to 5-HT inhibits pharyngeal pumping. Incubations in 300 μM tubocurarine had no inhibitory effect (n=8). **B.** Prolonged incubations of *G. rostochiensis* in 10 mM nicotine completely abolished 5-HT induced stylet thrusting. Effects of mecamylamine and tubocurarine on 5-HT induced stylet thrusting responses in significantly reduced (n=30). The data are shown as mean ± SEM and the analysis correspond to 5-HT as control or 5-HT + ligand and significance tested by one way ANOVA with Bonferroni’s multiple comparison (^ns^p>0.05, ****p<0.0001).

Chronic exposures of *G. rostochiensis* to nicotine, mecamylamine or tubocurarine, significantly improved their inhibitory effects on 5-HT stimulated stylet thrusting. A 24-hour incubation in nicotine prior to 5-HT exposure completely abolished stylet thrusting. Prolonged incubations in mecamylamine also inhibited the 5-HT induced stylet thrusting response. Interestingly, a prolonged incubation of *G. rostochiensis* in tubocurarine, which had no effect on 5-HT induced pharyngeal responses in *C. elegans* was quite potent. This significantly inhibited 5-HT induced stylet thrusting (**Fig. 5B**).

Tubocurarine has been reported to completely block ectopically expressed *Ce*.EAT-2 receptors at a concentration 10-fold lower than what was used in our whole animal assay. By breaking the cuticle of *C. elegans,* dissecting and exposing the pharyngeal muscles directly to tubocurarine, we significantly improved the drug potency (**Fig. 6A**). Mutants of EAT-2 which are physiologically defective in pharyngeal pumping were irresponsive to 5-HT stimulation and tubocurarine inhibition (**Fig. 6B**).

**Fig. 6.**
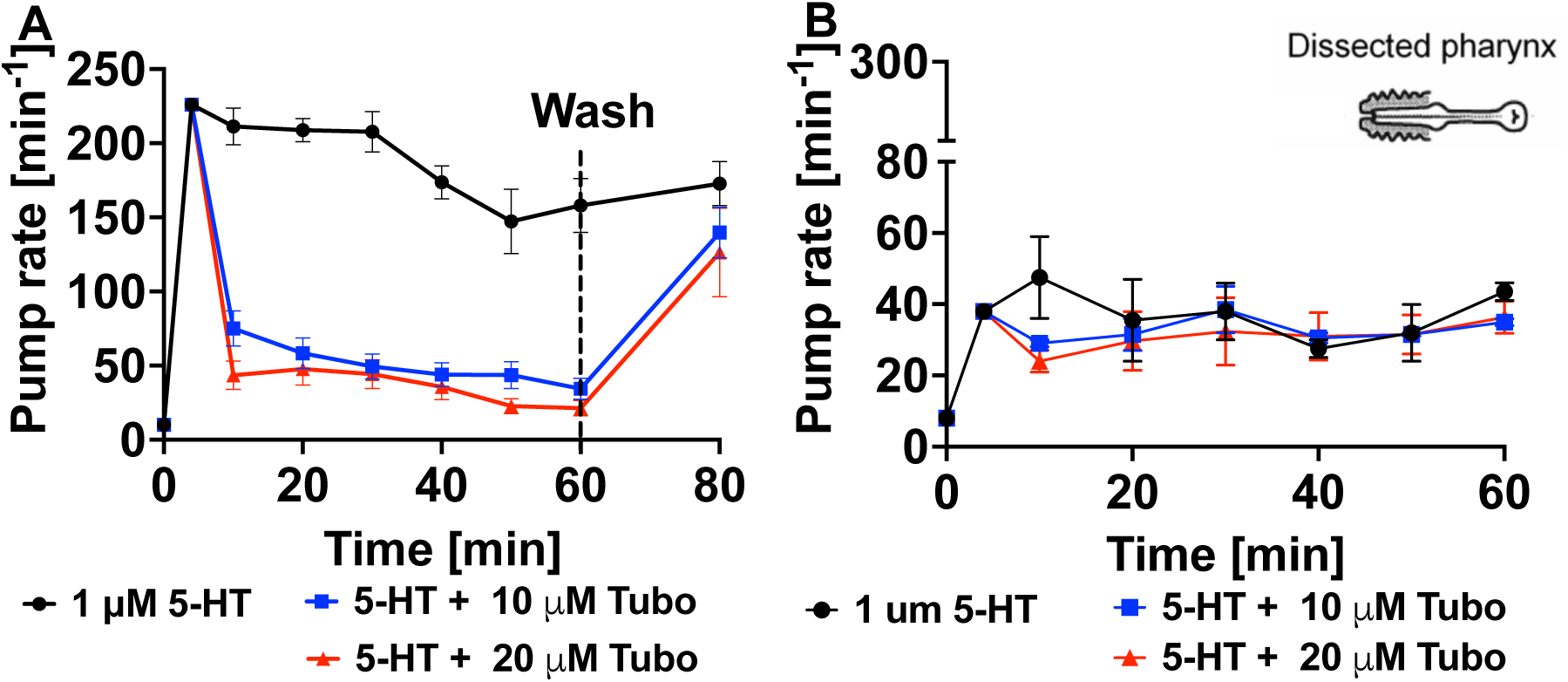
Inhibitory effect of tubocurarine on exposed *C. elegans* pharyngeal muscles. **A.** Tubocurarine inhibits 5-HT induced pharyngeal pumping in cut-head wildtype worms (N2). Pharyngeal pumping recovered after a 10-minute wash and re-exposure to 5-HT, n=10. **B.** Mutants of *eat-2(ad465)* show no response to 5-HT stimulation, n=3.

### Stylet thrusting and body posture in G. rostochiensis

As indicated above, in addition to inducing pharyngeal function the exogenous application of 5-HT induced changes in *G. rostochiensis* motility and body posture. When incubated for longer than 30 minutes in 1 mM 5-HT, G*. rostochiensis* became immobile and assumed a kink posture in the head region, with elevated stylet thrusting activity (**Fig. 7A**). However, *G. rostochiensis* incubated for 24 h in nicotine prior to 5-HT exposure failed to adopt this kinked posture and this coincided with the complete block of 5-HT induced thrusting. Interestingly, prolonged pre-incubations in mecamylamine and tubocurarine did not disrupt kinking but had a selective inhibitory effect on stylet thrusting (**Fig. 7B**). In summary, the comparison of the consequence of 5-HT on posture and stylet function in face of distinct nicotinic acetylcholine receptor agents suggests a potential uncoupling of the stylet thrusting and associated body postures.

**Fig. 7.**
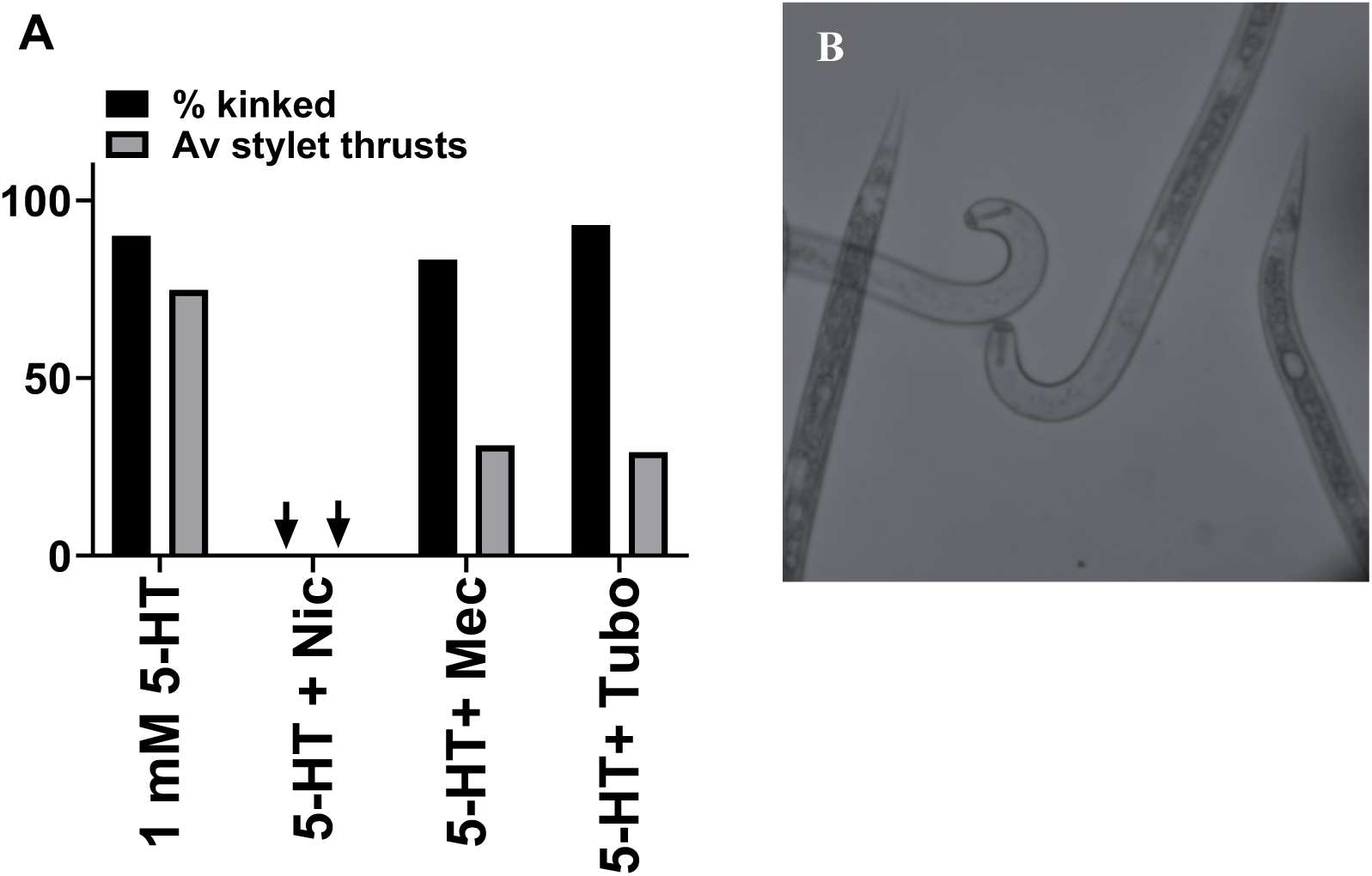
Comparing cholinergic drug effects on 5-HT induced stylet and posture states. **A.** The effects of 10 mM nicotine, 200 μM mecamylamine and 300 μM tubocurarine on 5-HT induced stylet thrusting and associated kinked posture on *G. rostochiensis* (5-HT n= 13, nicotine n=12, mecamylamine n=12 and tubocurarine n=30). J2s subjected to the indicated treatments were scored for their kinked body posture and stylet thrusting activity and represented as a percentage of kinked J2s and average (Av) stylet thrusts. **B.** Grayscale image of *G. rostochiensis* in a kinked posture when incubated in 1mM 5-HT.

## DISCUSSION

The targeting of several distinct molecules and functional loci of the cholinergic transmitter system has been a successful strategy for invertebrate pest control (Raisch and Raunser, 2023). These pesticides impact acetylcholinesterase, the vesicular acetylcholine transporter or distinct classes of acetylcholine receptors (Costa et al., 2008; Čadež et al., 2021; Goodchild et al., 2024) but their use is confounded by the off-target effects on other organisms (Wan et al., 2025). As a mitigation to this, nAChRs offered a more selective approach to inhibiting the nervous system of invertebrate pests (Bradford et al., 2020). However, despite phyla selective potency for this receptor class it is still challenging to overcome the issue of non-selective targeting of pests relative to non pest organisms (Li et al., 2025).

In this study we looked at specialized pharyngeal functions in two nematodes: stylet thrusting in the plant parasitic nematode *G. rostochiensis* and pharyngeal pumping in the free-living model nematode *C. elegans*. This could offer a tissue-selective target underpinning specialized life cycle behaviors as an approach to improve pesticide selectivity. We showed that these pharyngeal functions in both nematode species were controlled by cholinergic and serotonergic signaling. This is thought to be driven by the rather distinct, nematode-specific nAChR EAT-2 which has previously been shown to be required for normal pumping in *C. elegans* (McKay et al., 2004). It should be noted that whilst this role is important to *C. elegans* it is non-essential as null mutants are merely retarded in development (Avery, 1993; McKay et al., 2004). However, considering the multiplicity of life cycle selective functions like hatching, host root invasion and feeding behaviors that the pharyngeal muscles play in the PPNs it can be envisaged that targeting this receptor will be particularly pernicious for this nematode class. Our study probed the molecular and structural determinants of pharyngeal muscles that regulate pharyngeal function in *G. rostochiensis*. We investigated the possibility of translating existing understanding of EAT-2 function in *C. elegans* to PPNs.

Initially, we established EAT-2 presence in other nematode species through a protein blast and a phylogenetic compilation that revealed its conservation within Nematoda. This was confirmed in the PPN *G. rostochiensis* by successfully amplifying the cDNA for the predicted *Gr*.EAT-2 ORF. In *C. elegans*, EAT-2’s functional expression requires EAT-18, an auxiliary protein that is under investigated. EAT-18 is a short single transmembrane domain protein with an intracellular N-terminus and an extracellular C-terminus. Evidence suggests that in its absence, EAT-2 is made, trafficked and localized in the plasma membrane but is a non-functional receptor (Choudhary et al., 2020). This makes EAT-18 a potential target to indirectly disrupt EAT-2 function. We identified orthologues of *Ce*.EAT-18 in other nematode species, and a multiple sequence alignment of these orthologues revealed a high degree of conservation with a 59.09 % identity to the putative *Gr*.EAT-18 (see supplementary 2).

### Fluorescence in-situ hybridization reveals Gr.EAT-2 expression in the pharynx

Given the molecular conservation of EAT-2 across Nematoda, we probed its expression pattern and function. Such gene expression histology has been challenging in nematode species even with the increased availability of molecular information (Sperling and Eves-van den Akker, 2023). Our study took advantage of multiplex probes that facilitate the detection of gene expression in whole mount nematodes to provide primary evidence on the expression pattern of EAT-2 to underpin key signaling in stylet thrusting and possibly the median bulb pulsation. Most FISH studies on PPNs have focused on effector genes (De Boer et al., 1998, 1999; Lilley et al., 2018; Sperling and Eves-van den Akker, 2023). Here, we show visual evidence of the expression pattern for cholinergic neurons using probes for *Gr.unc-17,* the transporter gene required for loading synthesized ACh into vesicles (Alfonso et al., 1993), *Gr.eat-2*, a cholinergic receptor and putative regulator for pharyngeal function in PPNs and *Gr.myo-3* a highly expressed gene in longitudinal muscles of the body-wall. With probes for *Gr.eat-2* we identified discrete and reproducible expression in pharyngeal structures. The tissue in which we consistently found a hybridization pattern across several stained specimen was the median bulb (metacorpus). In addition, there was a robust but less frequent staining in discrete structures located in a position associated with the esophageal glands (**Fig. 2**). These observations resonate with expression patterns observed by McKay et al. (2004) and Cao et al. (2023). Sense probes for *Gr.unc-17* identified neurons in the pharynx and around the ventral side of the worm. The expression pattern for *Gr*.UNC-17 was similar to transgenic expression studies in *C. elegans* which showed *Ce*.UNC-17 expression in cholinergic neurons of the head region and the ventral nerve cord motor neuron (Mathews et al., 2012; Haque and Nazir, 2016). The selective distribution of cholinergic determinants is consistent with its key role in body wall muscle transmission, pharyngeal transmissions and supports a discrete role for EAT-2 in PPN pharyngeal function. Probes for *Gr.myo-3* localized along longitudinal muscles of the body wall muscle.

### A common pathway may regulate pharyngeal pumping and stylet thrusting

#### 5-HT indirectly regulates pharyngeal pumping and stylet thrusting

We proposed this molecular organization to investigate transmitter signaling in nematode pharyngeal function. Consistent with other findings, we showed that exogenous exposure to 5- HT induced the pharyngeal behaviors, pharyngeal pumping and stylet thrusting in *C. elegans* and *G. rostochiensis* respectively (Hobson et al., 2006; Horvitz et al., 1982; Perry et al., 2004). Although this happens, it is worth mentioning that 5-HT has no direct effect on EAT-2, the cholinergic regulator of pharyngeal pumping in *C. elegans.* The biogenic amine activates a cholinergic pathway through the MC neuron to initiate pharyngeal pumping via EAT-2 (McKay et al., 2004; Song and Avery, 2012). Beyond pharyngeal function, 5-HT concentrations that induce pharyngeal behaviors in *G. rostochiensis* rendered them immotile with a characteristic kinked-shape posture around the midbody or neck region. A supporting interpretation for this kink is that worms generate enough hydrostatic pressure and tension to act like a flex point for protractor muscles to drive stylet movements (Doncaster, 1966).

### ACh and nicotine directly induce feeding behaviors

The neuromuscular system within Nematoda shows a high degree of conservation, which allows for generally valid hypotheses and conclusions to be made on physiological behaviors like motility, egg laying, and feeding behaviors across different nematode species (Hahnel et al., 2020). *C. elegans* requires cholinergic signaling to achieve muscle contractions that drive pharyngeal functions like pharyngeal pumping and peristalsis. By utilizing the cholinergic compounds ACh and nicotine, we induced stimulatory effects on pharyngeal function with *C. elegans* and *G. rostochiensis*. Our findings complement the findings of Kozlova et al. (2019) who observed that WT *C. elegans* and *cha-1* mutants deficient in choline transferase activity (Rand and Russell, 1984) exposed to nicotine showed induced pharyngeal pumps whilst *eat-2* mutants were not significantly affected, suggesting that nicotine’s stimulatory effect on pharyngeal pumping may be EAT-2 dependent. ACh and nicotine have also been reported as agonist of recombinantly expressed EAT-2 receptors (Choudhary et al., 2020). Per se, comparing the pharmacologically induced pharyngeal behavior in *G. rostochiens* with the well-researched pharyngeal pump pathway in *C. elegans,* we propose that a cholinergic involvement via EAT-2 drives pharyngeal function that can be manifested as stylet thrusting (**Fig. 8**).

**Fig. 8.**
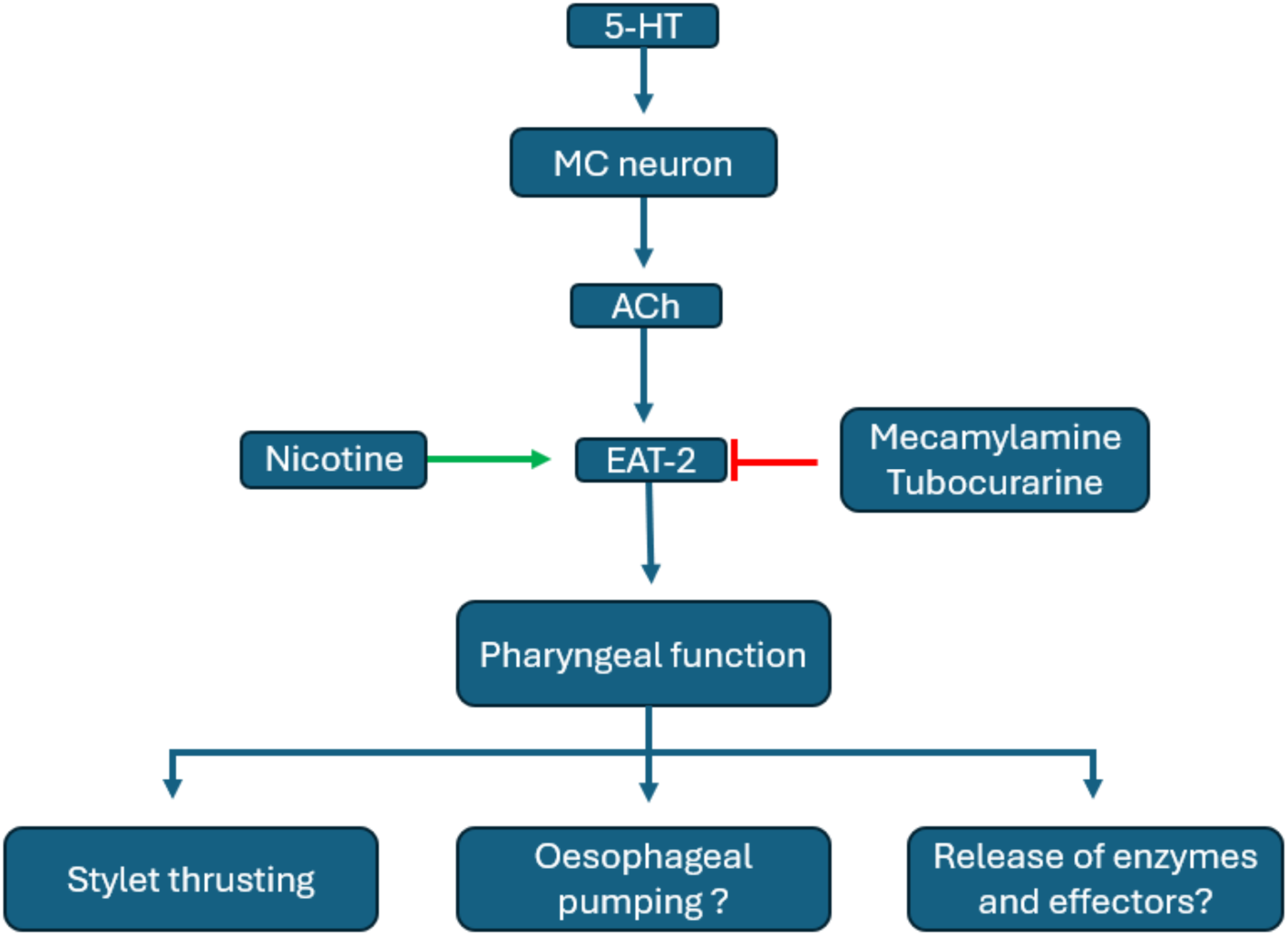
A putative signaling pathway for pharyngeal function in *G. rostochiesis*. 5-HT modulates pharyngeal function through the MC neuron, activating a cholinergic release that acts directly on EAT-2 and triggering pharyngeal function. Nicotine and ACh are agonists of EAT-2 that induce stylet thrusting. Mecamylamine and tubocurarine block EAT-2 and inhibit stylet thrusting functions.

The role of sensory inputs that innervate pharyngeal muscles and possibly controlling stylet protractor muscles have long been recognized (Doncaster, 1966). Here we provide insights into the downstream pharmacological and molecular determinants of this action.

### Nicotine inhibits 5-HT stimulated pharynx

Nicotine’s inhibitory effect on 5-HT stimulated pharyngeal responses was interesting because nicotine in solution induced pharyngeal responses but in combination with 5-HT was inhibitory, suggesting that the overstimulation of pharyngeal muscles resulted in an activation block and a subsequent inhibition in pharyngeal responses. Acute and chronic pre-exposures of *C. elegans* and *G. rostochiensis* to nicotine inhibit 5-HT induced pharyngeal effects and this may occur because the extended incubation leads to desensitization and an inhibitory block (Liu et al., 2025). This interpretation is in support of the notion that nicotinic responses act down stream of 5HT (Kudelska, 2019).

### Mecamylamine and tubocurarine directly pharyngeal function

We designed experiments to help decipher the intersection between pharyngeal pumping and stylet thrusting. The stimulatory effects of ACh and nicotine support the role of EAT-2 in these responses. Probing this further by investigating the effect of EAT-2 antagonists mecamylamine and tubocurarine on pharyngeal responses revealed their inhibitory significance on 5-HT stimulated pharyngeal responses. Thus, based on two chemically distinct inhibitors we can suggest that EAT-2 is an important mediator of stylet thrusting. The variation in drug potency from acute to chronic exposures and between intact worms and those whose cuticle had been broken open, reinforces the importance of the cuticle as a protective barrier (Johnstone, 1994). The differential drug effects observed from our assays between *C. elegans* and *G. rostochiensis* reflects on the differences in cuticle structure among nematode species (Decraemer and Hunt, 2013)

Our findings suggest that *Gr*.EAT-2 plays an important role in the signaling pathway that drives pharyngeal function and stylet thrusting. It will be interesting to understand how the discrete contexts that trigger stylet functions during the PPN lifestyle are integrated. Moreover, considering the biological significance of stylet thrusting for PPNs, we propose *Gr*.EAT-2 to be a valuable target to disrupt the lifecycle of this global economic agricultural pest. As such the EAT-2 pharmacophore merits further investigation to resolve potential selective channel modulators.

## Supporting information

https://sotonac.sharepoint.com/:f:/s/PhDdataNvenankeng/IgDMrH0S6l11Q7zHD40NVKDOAVuJk15fPfZxh8f-U-U5oFg?e=2XsMU2

