## Supplementary material for "Investigation of the nicotinic receptor EAT-2 as a novel target to mitigate plant parasitic nematode infections": https://sotonac.sharepoint.com/:f:/s/PhDdataNvenankeng/IgDMrH0S6l11Q7zHD40NVKDOAVuJk15fPfZxh8f-U-U5oFg?e=2XsMU2

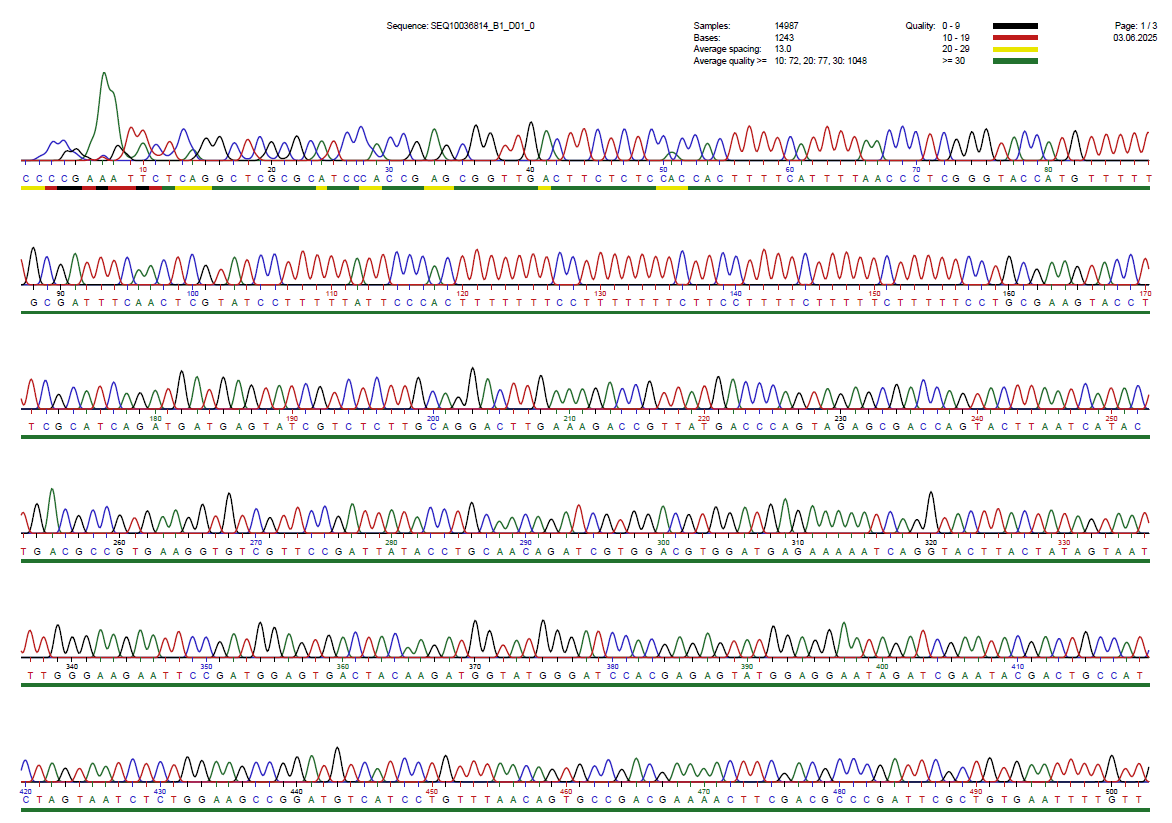

**A**

**START**

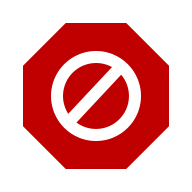

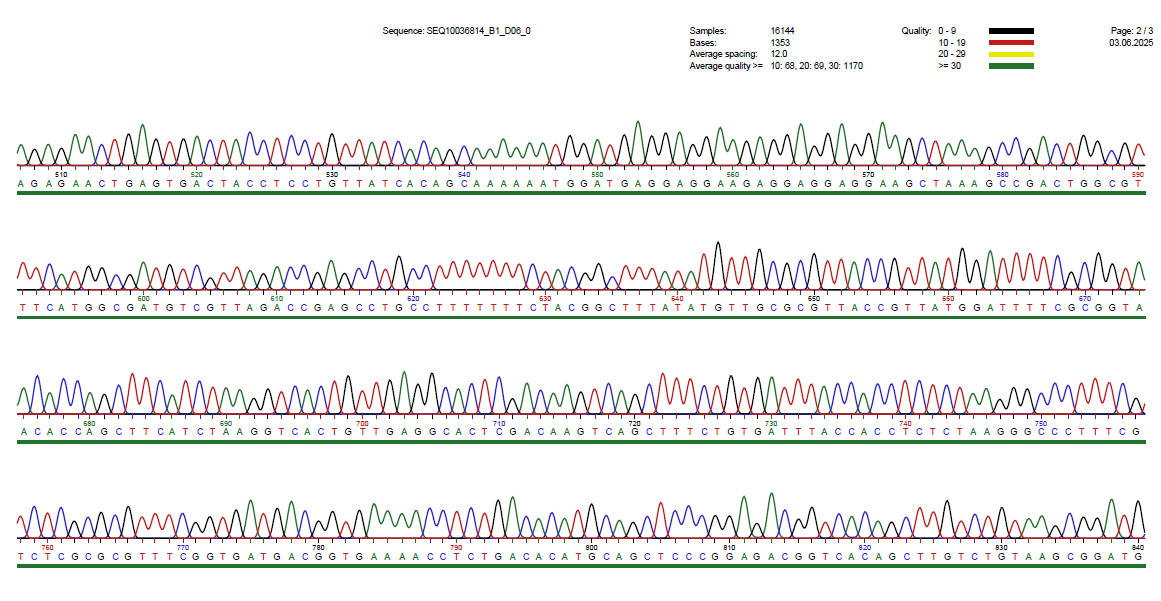

B

**STOP**

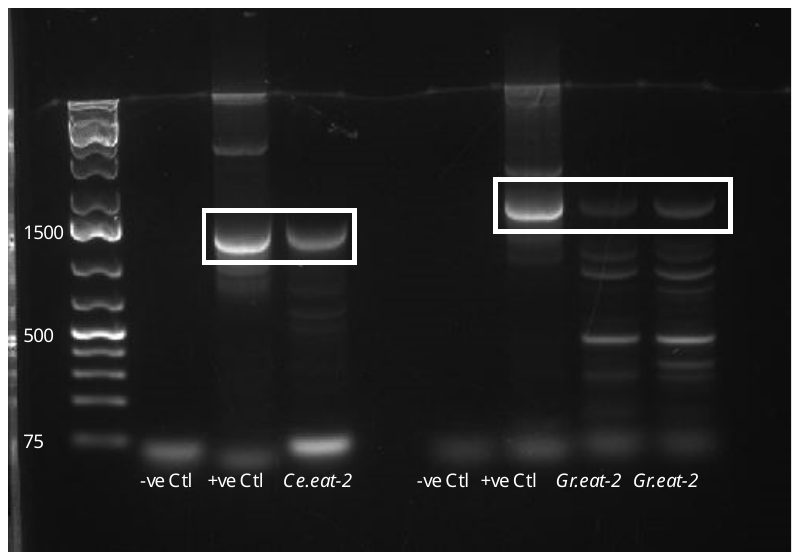

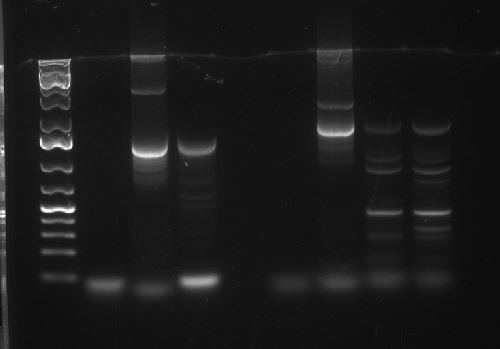

kb	1	2	3	4	5

**Supplementary 1. A.** Forward strand of PCR amplified *Gr*.EAT-2 transcript. Sequencing data reveals consistency with the predicted *Gr*.EAT ORF. **B.** Gel image of PCR amplified EAT-2 sequences for *C. elegans* and *G. rostochiensi* **1.** *Ce*.EAT amplified from a vector containing *Ce*.EAT-2*.* **2.** *Ce*.EAT-2 amplified from transcript isolated from ≈1K young adults. **3.** *Gr.*EAT-2 amplified from plasmids containing *Gr*.EAT-2 **4. & 5.** *Gr*.EAT-2 amplified from transcripts isolated from 2k and 8k J2s respectively.

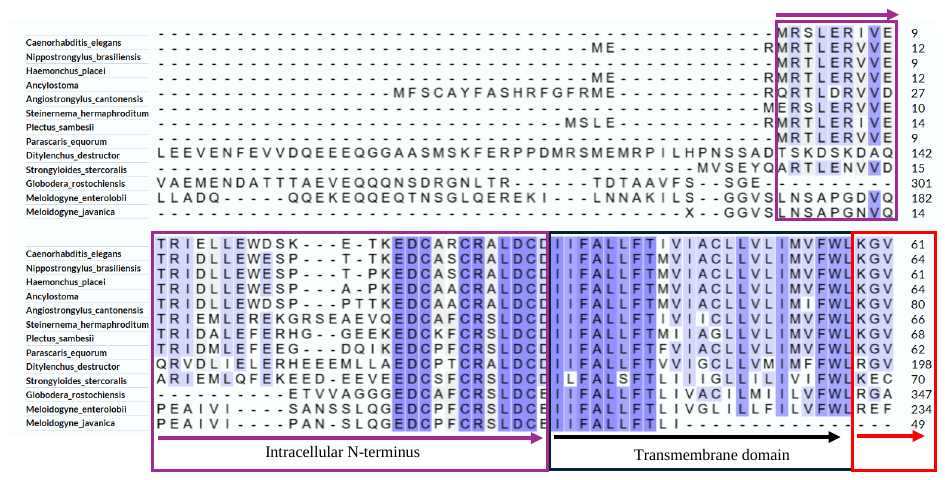

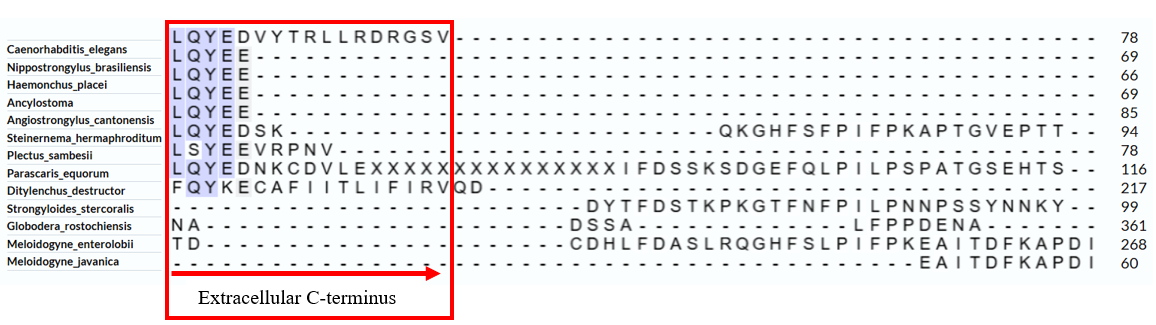

**Percentage identity**

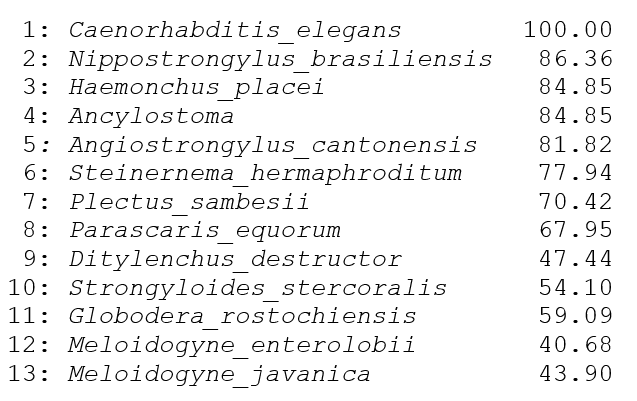

**Supplementary 2: EAT-18 transcript alignment for multiple nematode species.** The sequence alignment revealed a high degree of conservation of EAT-18 among free living, animal parasitic and plant parasitic nematodes with high identity percentages.
